# A meta-analysis of the effects of early life stress on the prefrontal cortex transcriptome suggests long-term effects on myelin

**DOI:** 10.1101/2024.11.22.624315

**Authors:** Toni Q. Duan, Megan H. Hagenauer, Elizabeth I. Flandreau, Anne Bader, Duy Manh Nguyen, Pamela M. Maras, Randriely Merscher S. De Lima, Trevonn Gyles, Christabel Mclain, Michael. J. Meaney, Eric J. Nestler, Stanley J. Watson, Huda Akil

## Abstract

**Background:** Early life stress (ELS) refers to exposure to negative childhood experiences, such as neglect, disaster, and physical, mental, or emotional abuse. ELS can permanently alter the brain, leading to cognitive impairment, increased sensitivity to future stressors, and mental health risks. The prefrontal cortex (PFC) is a key brain region implicated in the effects of ELS.

**Methods:** To better understand the effects of ELS on the PFC, we ran a meta-analysis of publicly available transcriptional profiling datasets. We identified five datasets (GSE89692, GSE116416, GSE14720, GSE153043, GSE124387) that characterized the long-term effects of multi-day postnatal ELS paradigms (maternal separation, limited nesting/bedding) in male and female laboratory rodents (rats, mice). The outcome variable was gene expression in the PFC later in adulthood as measured by microarray or RNA-Seq. To conduct the meta-analysis, preprocessed gene expression data were extracted from the Gemma database. Following quality control, the final sample size was n=89: n=42 controls & n=47 ELS: GSE116416 n=23 (no outliers); GSE116416 n=44 (2 outliers); GSE14720 n=7 (no outliers); GSE153043 n=9 (1 outlier), and GSE124387 n=6 (no outliers). Differential expression was calculated using the *limma* pipeline followed by an empirical Bayes correction. For each gene, a random effects meta-analysis model was then fit to the ELS vs. Control effect sizes (Log2 Fold Changes) from each study.

**Results:** Our meta-analysis yielded stable estimates for 11,885 genes, identifying five genes with differential expression following ELS (false discovery rate< 0.05): *transforming growth factor alpha* (*Tgfa*), *IQ motif containing GTPase activating protein 3* (*Iqgap3*), *collagen, type XI, alpha 1* (*Col11a1*), *claudin 11* (*Cldn11*) and *myelin associated glycoprotein* (*Mag*), all of which were downregulated.

Broadly, gene sets associated with oligodendrocyte differentiation, myelination, and brain development were downregulated following ELS. In contrast, genes previously shown to be upregulated in Major Depressive Disorder patients were upregulated following ELS.

**Conclusion:** These findings suggest that ELS during critical periods of development may produce long-term effects on the efficiency of transmission in the PFC and drive changes in gene expression similar to those underlying depression.

**Graphical Abstract:** 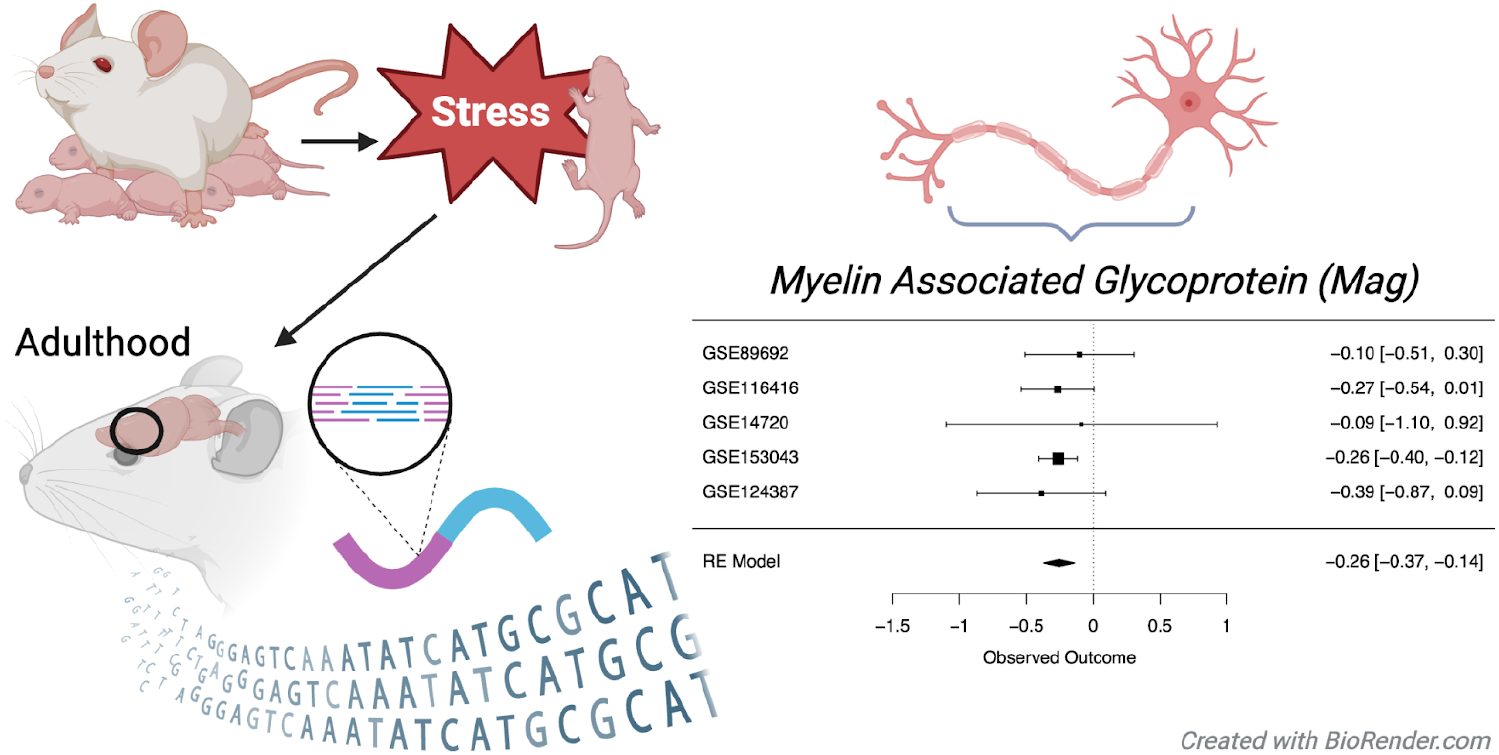

*Key Points:*

- Early life stress (ELS) can have long-term effects on the prefrontal cortex (PFC) and its related cognitive and emotional functions.
- To elucidate these long-term effects, we conducted a meta-analysis of five publicly available PFC transcriptional profiling datasets from adult rodents that had previously experienced ELS.
- This meta-analysis revealed a consistent downregulation of myelin-related genes in the PFC following ELS, and an upregulation of genes related to Major Depressive Disorder.

*Plain Language Summary:* Early life stress refers to exposure to negative childhood experiences, such as neglect, disaster, and physical, mental, or emotional abuse. Early life stress can permanently alter the brain, including the prefrontal cortex, which can lead to cognitive and emotional dysfunction that lasts into adulthood. We performed a meta-analysis using five public datasets to identify consistent long-term effects of early life stress on gene expression (mRNA) in the prefrontal cortex of adult rodents. In these studies, rodents that had experienced early life stress consistently showed a decreased amount of mRNA for genes related to myelin. Myelin is the fatty layer that insulates the axons of neurons, allowing them to transmit electrical signals more efficiently. These gene expression changes may suggest long-term effects of early life stress on the efficiency of prefrontal neurotransmission, disrupting cognitive and emotional processing.

## Introduction

Early life adversity, also known as early life stress (ELS), refers to exposure to negative childhood experiences. ELS can be caused by physical, mental, or emotional abuse, war, disaster, malnutrition, neglect, or negative parenting habits (Malave et al., 2022). Unfortunately, many children and adolescents around the world are exposed to ELS, with one in six children experiencing four or more adverse childhood experiences (Malave et al., 2022). These adverse experiences can have a long-lasting impact: studies have shown that ELS and adversity can permanently alter an individual’s brain (Barnett Burns et al., 2018), leading to cognitive impairment, sensitivity to future stressors, conduct disorder, mental health risks, diminished response to antidepressants, and other irreversible changes that persist into adulthood (Malave et al., 2022).

Researchers leverage animal models to study the effects of ELS on the brain. In laboratory settings, rodents typically reach significant developmental milestones within a short timeframe, progressing from birth to weaning in just three weeks (postnatal (P) days P0-P21). Although experiences throughout life are constantly altering the brain, the formation and functional maturation of the brain are shaped during this early postnatal period (Short & Baram, 2019). Circuits undergo critical periods during this time when certain experiences can lead to irreversible changes later in life (Short & Baram, 2019).

These periods of heightened plasticity are determined by neurobiological processes such as excitation and inhibition balance, synaptogenesis & synaptic pruning, myelination, and neurogenesis (Malave et al., 2022).

Much like humans, maternal care is critical for rodent development (Chen & Baram, 2016). Therefore, maternal separation is a widely studied animal model of ELS (Malave et al., 2022). During a maternal separation experiment, dams are separated from pups daily for 1 to 8 hours over a 2 to 3-week period. This separation typically starts 1 to 3 days postnatal. Maternal separation in rats increases offspring anxiety-like behaviors long-term. The effects of maternal separation on anxiety-like behaviors are more pronounced with a longer separation period (Malave et al., 2022), whereas a brief separation from the offspring, called early handling, can be protective (Pryce et al., 2005). The maternal privation model is the most extreme version of maternal separation. For this model, pups are removed from their mothers at 1-2 days postnatal and often fed using an artificial feeding device (Pryce et al., 2005).

A more recently developed rodent model of ELS is limited bedding and nesting material (Malave et al., 2022). In this model, rat or mouse dams and their pups are placed on a wire mesh with less standard nesting material and bedding material for the first week of life. This model replicates impoverished housing conditions by preventing the mother from building a suitable nest for pups. The lack of bedding and nesting conditions causes maternal care to be unpredictable, threats to recur, and abusive behavior to occur (Malave et al., 2022). The lack of bedding and nesting also reduces the pup’s body weight before weaning, which suggests malnutrition caused by deprivation (Gilles et al., 1996; Malave et al., 2022; Rice et al., 2008).

ELS is known to affect many key brain regions, including the hippocampus, anterior cingulate, ventral tegmental area, and prefrontal cortex (PFC) Peña et al., 2017, 2019; (Luo et al., 2023; Malave et al., 2022; Peña et al., 2017, 2019). ELS profoundly affects the structure and function of the anterior cingulate cortex, increasing the susceptibility to adult neuropsychiatric disorders, notably social dysfunction. Emotional and social brain networks centered on the PFC can be particularly affected by ELS, as can networks associated with executive functions and memory (Malave et al., 2022). Individuals with neuropsychiatric conditions have decreased hippocampal volume, especially if they experience stress before the age of thirteen (Malave et al., 2022). Evidence from rodent ELS models has similarly shown reduced hippocampal volume, reduced spine density, and impaired long-term potentiation (Malave et al., 2022).

To better understand the effects of ELS on the brain, we conducted a meta-analysis of publicly available transcriptional profiling datasets from rodent models. Transcriptional profiling technologies, such as RNA-Seq or microarray, provide information regarding which genes are active and quantify their expression levels in tissue samples, offering insights into biological processes, disease mechanisms, and responses to treatments. Despite the richness of this information, the conclusions drawn from individual transcriptional profiling studies are often limited due to small sample sizes and noise from technical variables. Meta-analysis is a useful tool for addressing these issues. By compiling data from several studies, meta-analysis can increase statistical power, identify consistent patterns across different datasets, and uncover more robust biological patterns. This allows us to derive more reliable conclusions and potentially discover novel associations that might not be evident in individual studies alone.

## Methods

### General

The meta-analysis pipeline was adapted from the *Brain Data Alchemy Project* (protocol: (M. Hagenauer et al., 2024), *not pre-registered*) and associated R code repository (https://github.com/hagenaue/BrainDataAlchemy). All analyses were conducted in R Studio using the programming language R (R v.4.2.1, R Studio v. 2023.06.0). The analysis code used for this project has been released at https://github.com/tonid2/EarlyLifeStressSummer2023.

To perform our meta-analysis of public transcriptional profiling data, we leveraged the data curation, preprocessing, and analysis efforts of the Gemma database (Lim et al., 2021). As of date, the Gemma project has reprocessed nearly 20,000 publicly available transcriptional profiling datasets.

Gemma’s preprocessing and data analysis steps adhere to a standardized pipeline. Gemma first realigns the datasets to an updated genome. Quality control measures performed by Gemma include identification and removal of outlier samples, removal of genes (rows) with minimal variance in expression values (either zero variance or <70% distinct values), and manual curation for common issues such as batch effects (Lim et al., 2021). Differential expression is calculated using the limma or limma-voom pipeline, and statistical output is available for the full model (omnibus) and individual contrasts.

### Dataset identification

The devtools R package (v.2.4.5, (Wickham et al., 2011) was used to install the *gemma.R* package from Github (gemma.R_1.3.2, https://github.com/PavlidisLab/gemma.R, (Lim et al., 2021; Zoubarev et al., 2012). The *search_datasets()* function was used to find transcriptional profiling datasets from laboratory rodents (rats or mice) within the Gemma database that contained prespecified keywords (**Figure 1**). We searched for datasets derived from three brain tissues implicated in the effects of ELS (regions of interest or ROIs): the hippocampus, anterior cingulate, and PFC. Later, we narrowed the focus of our meta-analysis to the PFC based on the number of available datasets.

**Figure 1.**
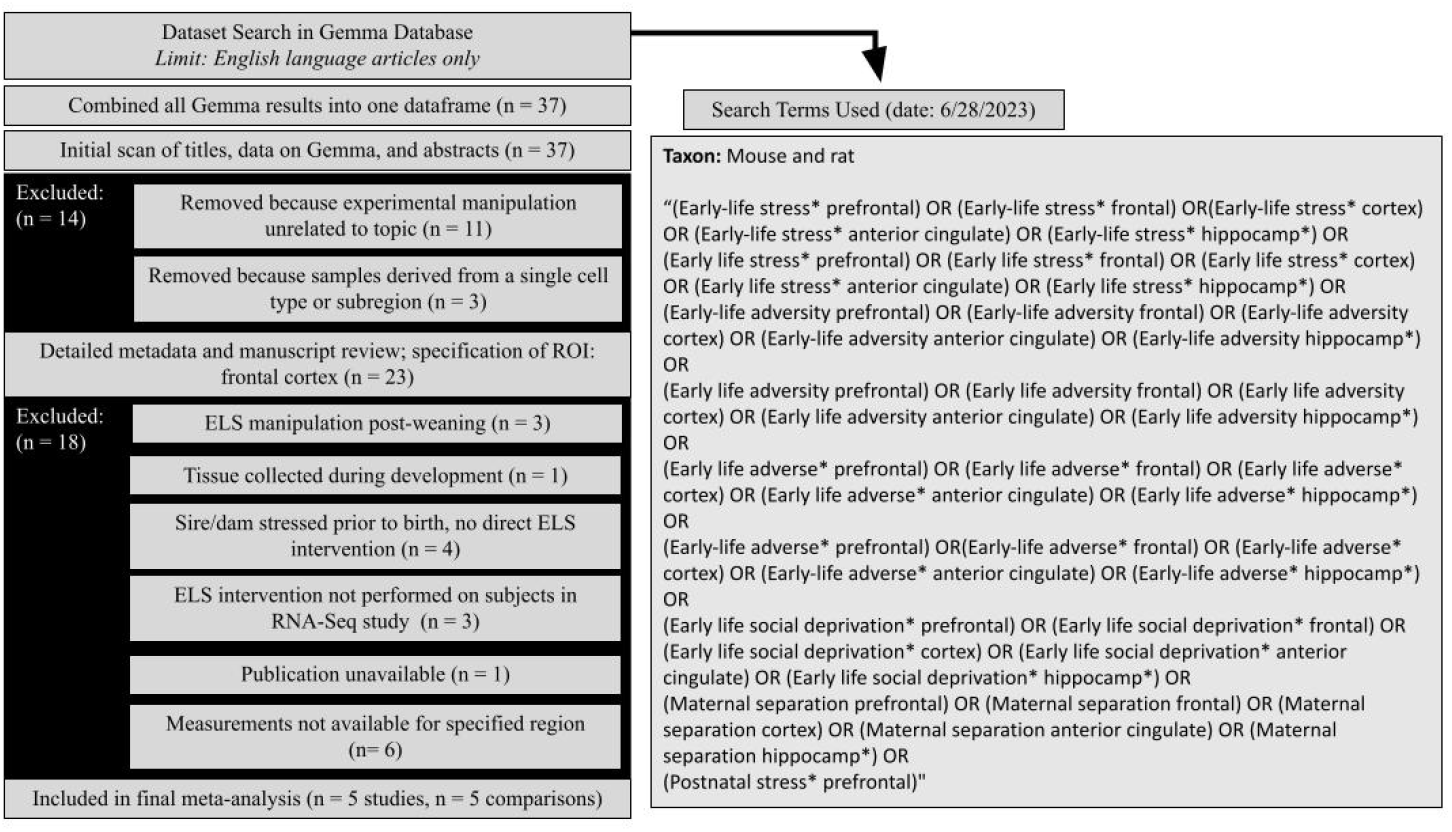
Diagram Overviewing Dataset Search and Selection. We identified transcriptional profiling datasets examining ELS models in laboratory rodents within the Gemma database using pre-specified keywords. Our initial search encompassed tissue from three brain regions implicated in the effects of ELS: the hippocampus, anterior cingulate, and prefrontal cortex (PFC). The focus of our meta-analysis was later narrowed to the prefrontal cortex based on the availability of datasets. The titles, abstracts, and metadata for the datasets were initially scanned and filtered using pre-specified inclusion/exclusion criteria, including indications that the dataset was not derived from a bulk dissection of brain tissue or that the content was unrelated to the research question (ELS). The secondary screening step included a detailed review of the metadata on Gemma and published methodology, followed by a final specification of the region of interest (ROI: PFC). Datasets were excluded in the secondary dataset filtering if they did not include an ELS manipulation during the pre-weaning period, the publication was not publicly unavailable, tissue was collected during development (instead of adulthood), or transcriptional profiling was not performed for the ROI. Abbreviations: n=number of datasets.

### Dataset filtering

The initial dataset filtering consisted of scanning dataset titles and metadata. The metadata records were filtered down to publicly available datasets, which were not labeled as problematic within the Gemma database. The titles, abstracts, and metadata for the datasets were scanned for immediate indications that the content was unrelated to our research question using pre-specified inclusion/exclusion criteria (protocol: (M. Hagenauer et al., 2024)) by a single researcher (TQD). Datasets were excluded if the experimental manipulation was unrelated to the meta-analysis topic (rodent models of ELS). Datasets were also excluded due to bulk brain tissue not having been collected in the experiment (*e.g*., scRNA-Seq, laser capture microscopy, cell sorting). After this filtering stage, a second researcher (MHH) approved all inclusion/exclusion choices.

The secondary dataset filtering involved specifying the brain ROI and then conducting detailed methodological reviews using both the available metadata and full-text manuscripts. As a brain ROI was not fully pre-specified in the search criteria, the research question was now narrowed based on dataset availability and filtered accordingly. For the three brain regions of interest, five available datasets represented the PFC, five datasets represented the hippocampus, and no datasets represented the anterior cingulate. The PFC was chosen as the final brain ROI.

The remaining dataset metadata records of interest were then subjected to a detailed review of accompanying experimental methods and available file formats. For this level of filtering, the publications associated with the datasets were referenced in addition to the detailed metadata available on Gemma. Only the methodological information in the publication was used to determine inclusion/exclusion. When reviewing the methodological information in the publications, we used a set of pre-specified inclusion/exclusion criteria (M. Hagenauer et al., 2024), as well as criteria specific to our research question, including the requirement that the ELS occur during the postnatal pre-weaning period and tissue be collected in adulthood. One experiment included a prenatal immune challenge (lipopolysaccharide on embryonic day 17) in addition to ELS. Otherwise, the risk of systematic bias within the datasets from the individual transcriptional profiling studies was deemed low for any particular gene-level result, as the original publications conducted full genome analyses. At this stage, all inclusion/exclusion decisions were reviewed by the entire Summer 2023 cohort in the Brain Data Alchemy Project (MHH, EIF, AB, TG, and CM).

### Overview of the final included datasets

After initial and secondary dataset filtering, a total of five datasets fulfilled all requirements and were used in the meta-analysis (**Table 1**: GSE89692, GSE116416, GSE14720, GSE153043, GSE124387 - *please note that GSE124387 was initially accidentally miscategorized as an exclusion*). Out of the five, three datasets were measured by RNA-seq, while two were measured by microarray. The ELS paradigms included some combination of maternal separation for >3 hours/day or limited nesting and bedding in male and female rodents (mice or rats). The rodents were stressed during pre-weaning, within time periods ranging from postnatal day 1 (P1) to day 21 (P21). In all five studies, PFC tissue was collected later during adulthood; two studies specifically sampled the medial PFC.

**Table 1.** Overview of Studies Included in the Meta-Analysis. Within the table, “GEO ID #” refers to the Gene Expression Omnibus (GEO) accession number for the dataset. The sample size (“n”) for the full dataset is provided, as well as the final sample size for the control (“Ref”) and ELS groups included the subset of the dataset used for our analysis (final total n: Ref=42, ELS=47). The subsetted sample size only includes the ROI (prefrontal cortex (PFC) or medial PFC (mPFC)) and excludes outlier samples. Please note that each sample in GSE89692 represents RNA pooled from 3 subjects. The Author and Year refer to the publication (Benekareddy et al., 2010; Oldham Green et al., 2021; Peña et al., 2017; Rincel et al., 2019; Zheng et al., 2019). “Duration” indicates how long (hours: “hrs”) the pups experienced maternal separation each day; limited nesting and bedding lasted all day. The “P” under “Age” represents the postnatal days during which the rodents were stressed. In GSE116416, the ELS group also received a prenatal immune challenge of lipopolysaccharide (LPS) on embryonic day 17 (E17). “Age Tissue” indicates the age of sacrifice, when the tissue from the ROI was collected. All experimental details refer specifically to the samples used in the transcriptional profiling experiment and not the full paper.

| GEO ID # | n (Full Dataset) | # Outliers | Final n (Ref) | Final n (ELS) | Species | Author | Year | Platform | Tissue | Type | Duration | Age Stressed | Age Tissue | Sex |
| --- | --- | --- | --- | --- | --- | --- | --- | --- | --- | --- | --- | --- | --- | --- |
| GSE89692 | 163* | 0 | 12 | 11 | Mouse | Peña et al. | 2017, 2019 | Illumina (RNA-seq) | mPFC | Maternal Separation & Limited Nesting | 4 hrs/day Separation | P10 - 20 (Male); P10 - 17 (Female) | Adult | Both |
| GSE116416 | 47 | 2 | 20 | 24 | Mouse | Rincel et al. | 2019 | Agilent (Microarray) | mPFC | Maternal Separation, Maternal Stress, & Prenatal LPS | 3 hrs/day Separation | P2 - P14; Prenatal LPS: E17 | Adult | Both |
| GSE14720 | 11 | 0 | 3 | 4 | Rat | Benekareddy et al. | 2010 | Agilent (Microarray) | PFC | Maternal Separation | 3 hrs/day Separation | P2 - P14 | Adult | Male |
| GSE153043 | 10 | 1 | 4 | 5 | Rat | Green et al. | 2021 | Illumina (RNA-seq) | PFC | Limited Nesting | Full-day | P2 - P9 | Adult | Male |
| GSE124387 | 9 | 0 | 3 | 3 | Rat | Zheng et al. | 2019 | Illumina<br>(RNA-seq) | PFC | Maternal<br>Separation | 4 hrs/day<br>Separation | P1 - P21 | Adult | Male |

### Individual dataset reprocessing: Outliers, gene filtering, and batch effects (quality control)

When possible, the preprocessed gene expression data for the selected experiments were re-analyzed to double-check data quality and ensure preferred sample subsetting and differential expression model specification (GSE89692, GSE116416, GSE14720, GSE153043). To do this reanalysis, the summarized experiment objects for each study were read into R using *Gemma.R* to access the API for the Gemma database (July 2023). For one dataset, the preprocessed gene expression data (summarized experiment object) were not available (GSE124387). For this dataset, we reviewed the data quality and model specification provided by Gemma, and have provided that information below.

Each of the summarized experiment objects was subsetted to the relevant samples for the meta-analysis. We subsetted by “organism part” (to focus on the PFC) and “treatment” (to only include the ELS and reference subject role subjects). Gemma defines outliers as samples with an adjusted median sample-sample correlation that is outside the interquartile range of the sample-sample correlations for the sample with the closest median correlation to them (Lim et al., 2021). We double-checked whether individual samples, as well as pairs or groups of outliers, might meet similar criteria regarding distance from the rest of the dataset by visualizing the distribution of sample-sample correlations within each dataset using a correlation matrix and boxplots. After visualization of heat maps and boxplots, one outlier was removed from dataset GSE153043, while two outliers were removed from dataset GSE116416 (**Table 1**).

Within the Gemma database, batch-related variables are combined into a single column (“block”). We used *strsplit()* to divide “block” into specific batch-related variables. For RNA-Seq, these variables included scan date, device, and run, which often impact gene expression measurements, as well as lane and flow cell. To determine if any of these batch-related variables might confound our design, we created cross-tables comparing each batch-related variable to our main variables of interest. We also examined the relationship between library size and our variables of interest using boxplots. To determine the impact of batch-related variables on our datasets, we used principal components analysis (PCA) to identify the main patterns of variation within gene expression data using the *prcomp(x, scale=TRUE)* function from the *stats* package, as applied to the transposed log2 gene expression matrix. The top four principal components of variation (PC1-4) were compared to our variables of interest and potential nuisance variables (batch, library size, and other experimental factors in the dataset) using boxplots and scatterplots. This information was used to choose the covariates included in the differential expression model for each dataset.

Only one dataset ended up having a differential expression model that included covariates (GSE89692, **Eq.1**). The differential expression models for the other datasets were kept simple **(Eq.2**), as these datasets were either very small (GSE153043, GSE14720 & GSE124387) or processed in a single batch (GSE116416) and contained samples with homogenous features within the specified subset.

**Eq.1 The differential expression model for GSE89692:**

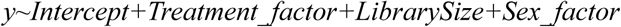

**Eq.2 The differential expression model for GSE116416, GSE153043, GSE14720 & GSE124387:**

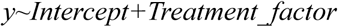

To mirror Gemma’s pipeline, we calculated ELS vs. Control differential expression using the *limma* pipeline, accounting for the mean-variance relationship present in the datasets (*limma-trend*, GSE124387: *limma-voom*), and then applied an empirical Bayes correction to the differential expression results using the *eBayes()* function (Ritchie et al., 2015).

### Result extraction

The result extraction process was similar to the Brain Data Alchemy Pipeline Methods for Summer 2023 (M. Hagenauer et al., 2024). Using either our own reprocessed differential expression results (GSE89692, GSE116416, GSE14720, GSE153043) or the differential expression results provided by Gemma (GSE124387), we extracted the Log2 Fold Changes (Log2FC) and sampling variances from the ELS vs. control differential expression results for each dataset. To be included in the meta-analysis, a gene needed to be present in the results from a minimum of four out of the five datasets. 11,889 genes fulfilled this criterion, and 11,885 genes produced stable meta-analysis estimates using a simple (intercept only) random effects meta-analysis model within the *metafor* package (Viechtbauer, 2010). The final sample size was not deemed large enough to conduct additional analyses to assess dataset heterogeneity or the effects of potential variables of influence (*e.g*., ELS type). The meta-analysis p-values then underwent false discovery rate (FDR) correction using the Benjamini-Hochberg method, also known as “q-value,” implemented via the *mt.rawp2adjp(x, proc=c(“BH”))* function in the *multtest* package (v.2.8.0, Pollard et al., 2005). Statistical significance was defined at a 5% FDR threshold (FDR<0.05). To highlight key findings, forest plots depicting effect sizes (Log2FC) and their 95% confidence intervals from each contributing study were generated using the *forest.rma()* function from the *metafor* package.

### Exploring functional patterns within the meta-analysis results

*StringDB:* StringDB was used to explore potential protein–protein interactions between the top differentially expressed genes identified by the meta-analysis. The StringDB database contains predicted protein-protein interaction (PPI) information from numerous sources, including experimental data, computational prediction methods, and public text collections (https://string-db.org/, accessed on September 27, 2024 (Szklarczyk et al., 2023)). The mouse gene symbols for the top genes (p<0.001: 40 genes) from our meta-analysis were entered into the StringDB database (minimum required interaction score: medium confidence (0.40)), and *Mus musculus* was selected as the species of interest. When examining the functional gene sets enriched within our network, we used a statistical background of all 11,889 genes included in our meta-analysis.

*Brain.GMT:* Brain.GMT is a curated database of 918 gene sets related to nervous system function, tissue, and cell types. A version of Brain.GMT that has been combined with a more traditional gene ontology gene set database (MSigDB “C5”: 14,996 gene sets) can be used within common analysis pipelines (GSEA, limma, edgeR) to interpret neuroscience transcriptional profiling results from three species: rat, mouse, and human. Brain.GMT can substantially improve the interpretation of differential expression results (M. H. Hagenauer et al., 2024). We adapted the example R code provided with the release of Brain.GMT that illustrates the use of Brain.GMT within a Fast Gene Set Enrichment Analysis. Prior to running the code, gene sets specifically derived from other brain tissues (the hippocampus or nucleus accumbens, as marked in Table 1 of the Brain.GMT paper: (M. H. Hagenauer et al., 2024)) were trimmed out of the .GMT file. Additionally, any cell type gene sets (as marked “C8” in Table 1 of the Brain.GMT paper (M. Hagenauer et al., 2024)) that were from tissues that were highly distinct from cortex were trimmed out, with the exception of bone marrow, which can reflect blood cell types (as there is blood in brain tissue).

## Results

To be included in the meta-analysis, a gene needed to be present in the results from a minimum of four out of five datasets. 11,889 genes fulfilled this criterion, and 11,885 genes produced stable meta-analysis estimates (**Table S1**). Of the 11,885 genes represented in the meta-analysis, five genes were significantly differentially expressed (FDR<0.05) in ELS models (**Table 2**): *transforming growth factor alpha* (*Tgfa*, **Figure S1A**), *claudin 11* (*Cldn11*, **Figure 2A**), *myelin-associated glycoprotein* (*Mag*, **Figure 2B**), *collagen, type XI, alpha 1* (*Col11a1*, **Figure S1B**), and *IQ motif containing GTPase activating protein 3* (*Iqgap3*, **Figure S1C**). All were down-regulated. Twelve other genes showed a trend towards being differentially expressed (FDR<0.10), with six of those genes down-regulated by stress (**Table 2**) and six genes upregulated by stress (**Table 2**).

**Table 2.**
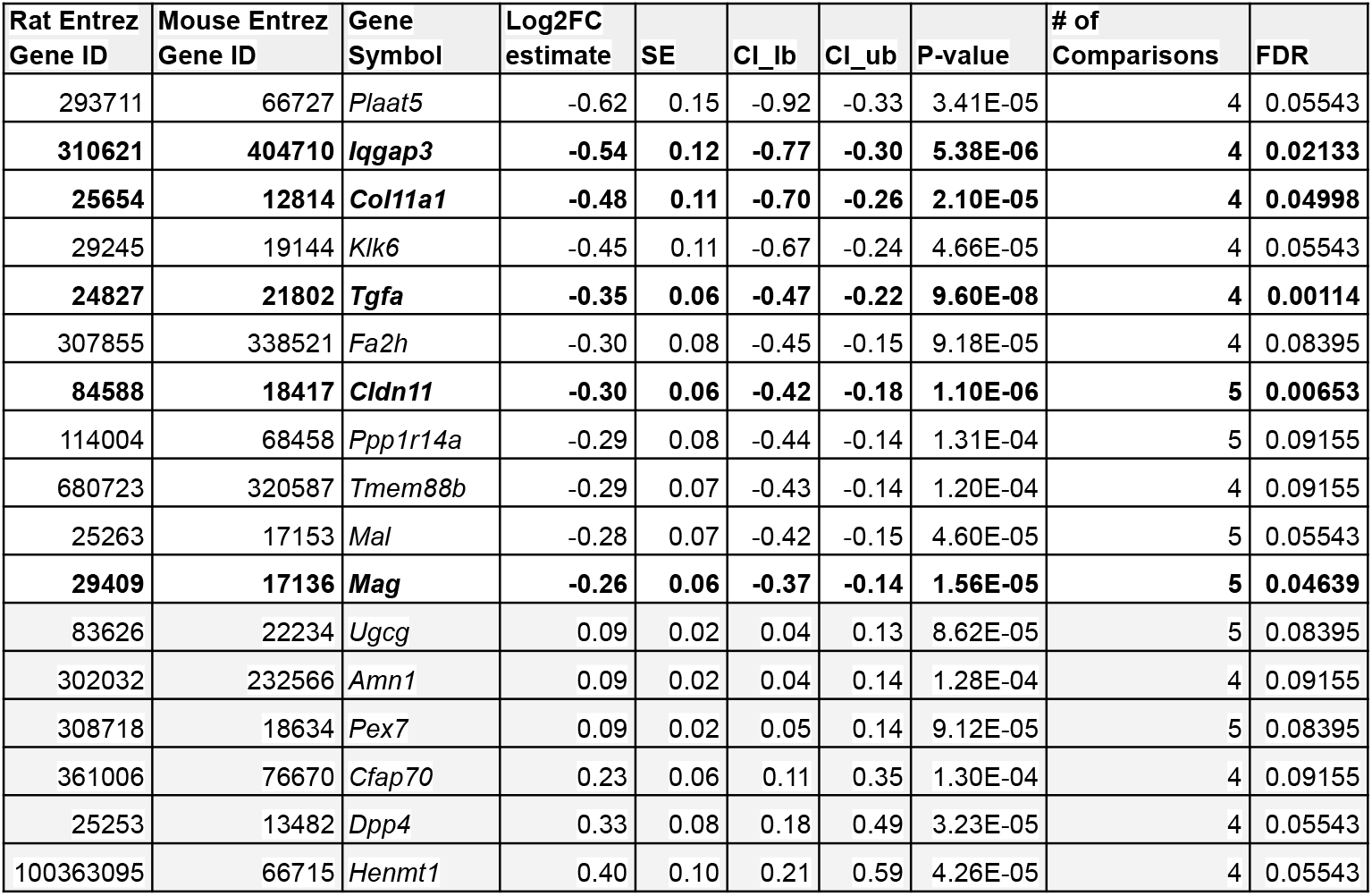
Meta-Analysis Results: Top Differentially Expressed Genes (FDR<0.10) in the PFC of Adult Rodents That Had Experienced ELS. Rows highlighted with bold text denote genes that are significantly differentially expressed (FDR<0.05), all other genes show a trend towards differential expression (FDR<0.10). The rows are ordered by Log2FC (lowest to highest). Definitions: Log2FC = Log2 Fold Change (ELS vs. Control), SE = Standard Error, CI_lb=confidence interval lower bound, CI_ub=confidence interval upper bound, p-value = nominal p-value, FDR = False Discovery Rate (q-value). A positive Log2FC denotes that a gene’s expression is higher in the experimental group compared to the control group (upregulated: highlighted grey), with higher positive values indicating greater increases. A negative Log2FC indicates lower expression in the experimental group relative to the control (downregulated), where larger negative values signify more substantial decreases. The # of comparisons represents the number of datasets that included measurements for the gene following quality control.

| Rat Entrez Gene ID | Mouse Entrez Gene ID | Gene Symbol | Log2FC estimate | SE | CI_lb | CI_ub | P-value | # of Comparisons | FDR |
| --- | --- | --- | --- | --- | --- | --- | --- | --- | --- |
| 293711 | 66727 | *Plaat5* | -0.62 | 0.15 | -0.92 | -0.33 | 3.41E-05 | 4 | 0.05543 |
| **310621** | **404710** | ***Iqgap3*** | **-0.54** | **0.12** | **-0.77** | **-0.30** | **5.38E-06** | **4** | **0.02133** |
| **25654** | **12814** | ***Col11a1*** | **-0.48** | **0.11** | **-0.70** | **-0.26** | **2.10E-05** | **4** | **0.04998** |
| 29245 | 19144 | *Klk6* | -0.45 | 0.11 | -0.67 | -0.24 | 4.66E-05 | 4 | 0.05543 |
| **24827** | **21802** | ***Tgfa*** | **-0.35** | **0.06** | **-0.47** | **-0.22** | **9.60E-08** | **4** | **0.00114** |
| 307855 | 338521 | *Fa2h* | -0.30 | 0.08 | -0.45 | -0.15 | 9.18E-05 | 4 | 0.08395 |
| **84588** | **18417** | ***Cldn11*** | **-0.30** | **0.06** | **-0.42** | **-0.18** | **1.10E-06** | **5** | **0.00653** |
| 114004 | 68458 | *Ppp1r14a* | -0.29 | 0.08 | -0.44 | -0.14 | 1.31E-04 | 5 | 0.09155 |
| 680723 | 320587 | *Tmem88b* | -0.29 | 0.07 | -0.43 | -0.14 | 1.20E-04 | 4 | 0.09155 |
| 25263 | 17153 | *Mal* | -0.28 | 0.07 | -0.42 | -0.15 | 4.60E-05 | 5 | 0.05543 |
| **29409** | **17136** | ***Mag*** | **-0.26** | **0.06** | **-0.37** | **-0.14** | **1.56E-05** | **5** | **0.04639** |
| 83626 | 22234 | *Ugcg* | 0.09 | 0.02 | 0.04 | 0.13 | 8.62E-05 | 5 | 0.08395 |
| 302032 | 232566 | *Amn1* | 0.09 | 0.02 | 0.04 | 0.14 | 1.28E-04 | 4 | 0.09155 |
| 308718 | 18634 | *Pex7* | 0.09 | 0.02 | 0.05 | 0.14 | 9.12E-05 | 5 | 0.08395 |
| 361006 | 76670 | *Cfap70* | 0.23 | 0.06 | 0.11 | 0.35 | 1.30E-04 | 4 | 0.09155 |
| 25253 | 13482 | *Dpp4* | 0.33 | 0.08 | 0.18 | 0.49 | 3.23E-05 | 4 | 0.05543 |
| 100363095 | 66715 | *Henmt1* | 0.40 | 0.10 | 0.21 | 0.59 | 4.26E-05 | 4 | 0.05543 |

**Figure 2.**
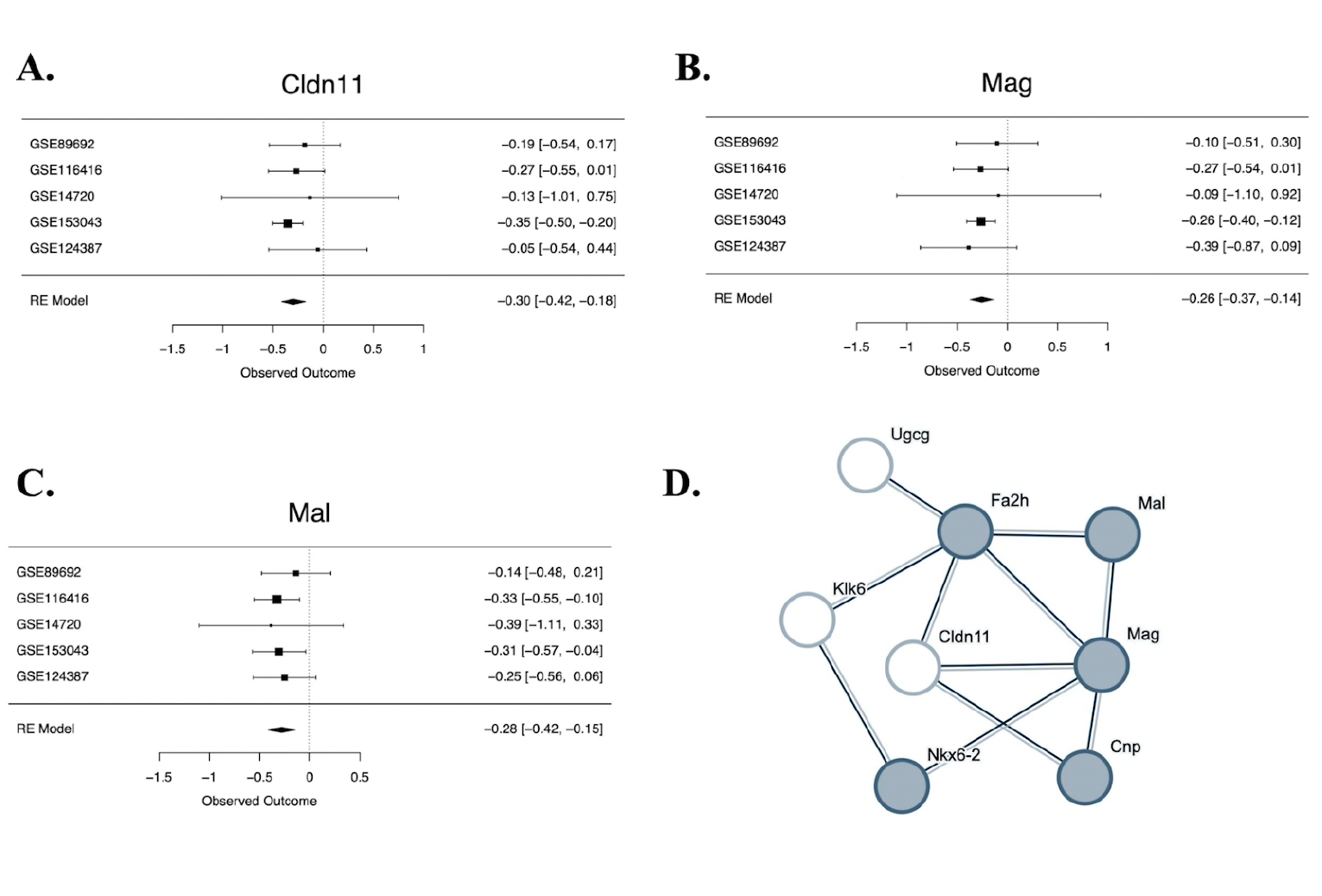
Genes within a network focused on oligodendrocyte differentiation are differentially expressed in early life stress models. **A-C**. Forest plots for two differentially expressed genes from the meta-analysis (FDR<0.05: Cldn11 & Mag), and a third gene that shows a trend towards differential expression (FDR<0.10: Mal) that are all part of a network related to oligodendrocyte differentiation. Rows illustrate ELS Log2FC (squares) with 95% confidence intervals (whiskers) for each of the datasets and the meta-analysis random effects model (“RE Model”). Forest plots allow for visual inspection of the consistency and magnitude of effects across the five studies. **A**. A forest plot showing the down-regulation of claudin 11(Cldn11) in ELS models. **B**. A forest plot showing the down-regulation of myelin-associated glycoprotein (Mag) in ELS models. **C**. A forest plot showing the trend towards a down-regulation of mal, T cell differentiation protein (Mal) in early life stress models. **D**. Multiple top differentially expressed genes are found in the same predicted protein-protein interaction (PPI) network enriched for genes associated with oligodendrocyte differentiation. To create the PPI network, top genes from the meta-analysis (p<0.001: 40 genes) were entered into the StringDB database. The only identifiable network (cluster) of 8 genes included six genes with either significant differential expression in ELS models (FDR<0.05: Mag, Cldn11) or a non-significant trend towards differential expression in ELS models (FDR<0.10: Mal, Fa2h, Klk6, Ugcg). The nodes represent proteins and are labeled with mouse gene symbol annotation. The lines represent predicted protein-protein associations. The associations are meant to be specific and meaningful (proteins jointly contribute to a shared function), but this does not necessarily mean they are physically binding to each other. The shaded nodes (grey) were identified as being part of a gene set involved in oligodendrocyte differentiation (GO:0048709) that was enriched with differential expression. Other genes within the network are part of a gene set involved in Central nervous system myelination (GO:0022010) that was also enriched with differential expression (including Klk6), or have also been previously associated with oligodendrocyte function and myelination (Cldn11).

To explore the biological functions associated with our differentially expressed genes, we first created a predicted PPI network by entering top genes from the meta-analysis (p<0.001: 40 genes) into the StringDB database. A network (cluster) of 8 genes was identified that included several significantly differentially expressed genes (FDR<0.05: *Cldn11, Mag*) (**Figure 2D**). To identify functional gene sets enriched within our PPI network, we compared the top genes (p<0.001: 40 genes) from the meta-analysis to a statistical background of all of the 11,889 genes included in the meta-analysis. Five of the genes that were present in the PPI network were identified as involved in *oligodendrocyte differentiation (GO:0048709)*. In contrast, only 85 proteins in total (in the network and the background) have this term assigned. This showed that there was an enrichment of overlap with our ELS network (FDR=0.0488, enrichment=1.24). Four genes within the PPI network were also identified as involved in the related concept of *Central nervous system myelination (GO:0022010)*, with only 27 proteins in total (in the network and the background) having this term assigned (FDR=0.0398, enrichment=1.64).

For a more comprehensive identification of functional gene sets enriched with the effects of ELS in our meta-analysis, we performed a fast gene set enrichment analysis (fGSEA) using both a brain-related functional gene set database (*Brain.GMT*) and traditional gene ontology gene sets. In the gene set enrichment analysis, we found that 42 gene sets (out of 9322 gene sets included in the results) had an enrichment with the effects of ELS that survived FDR correction (FDR<0.05, **Table S2**). Of these 42 gene sets, 21 gene sets were from *Brain.GMT* and 21 were traditional gene ontology gene sets. Most of these enriched gene sets were down-regulated following ELS (40 of 42), and explicitly related to oligodendrocyte development or myelination (22 of 42), or more broadly related to brain development (6 of 42). There were two notable exceptions: one down-regulated gene set was specifically related to stress hormone signaling (*GOBP_RESPONSE_TO_MINERALOCORTICOID*, strongest leading gene: *neuronal PAS domain protein 4* (*Npas4*), Log2FC=-0.34, p=0.012, FDR=0.61), and one of the only upregulated gene sets was derived from a meta-analysis characterizing the effects of Major Depressive Disorder in the frontal cortex (*Gandal_2018_MajorDepressiveDisorder_Upregulated_Cortex*, strongest leading gene: *follistatin* (*Fst*): Log2FC=0.14, p=0.0092, FDR=0.60). Twenty five of the enriched gene sets contained leading genes that were significantly differentially expressed genes following ELS in the meta-analysis (*Tgfa, Cldn11, Mag, Col11a1;* listed in **Table 3**).

**Table 3.** Functional gene sets that are enriched with the effects of early life stress are often related to oligodendrocyte development and myelination. Out of 42 gene sets that were enriched with the effects of ELS (FDR<0.05) within a gene set enrichment analysis performed using the meta-analysis results, most were down-regulated following ELS (40 of 42), and explicitly related to oligodendrocyte development or myelination (22 of 42). Listed above are the 25 enriched gene sets that contained leading genes that were significantly differentially expressed genes following ELS in the meta-analysis (Tgfa, Cldn11, Mag, Col11a1). P-value: Nominal p-value, FDR: False discovery rate, ES: Enrichment Score, NES: Normalized Enrichment Score, Size: Size of gene set, “Leading Genes Include”: a list of the leading genes in the gene set that were significantly differentially expressed following ELS (FDR<0.05) in the meta-analysis.

| Gene Set | P-value | FDR | ES | NES | Size | Leading Genes Include: |
| --- | --- | --- | --- | --- | --- | --- |
| GOMF_SIGNALING_RECEPTOR_BINDING | 0.00015 | 0.045 | -0.31 | -1.38 | 959 | *Tgfa*, *Mag* |
| GOBP_BIOLOGICAL_ADHESION | 0.00015 | 0.045 | -0.31 | -1.39 | 967 | *Cldn11*, *Mag* |
| GOBERT_OLIGODENDROCYTE_DIFFERENTIATION_DN | 0.00015 | 0.045 | -0.37 | -1.64 | 891 | *Col11a1* |
| GOBP_NEURON_DEVELOPMENT | 0.00015 | 0.045 | -0.33 | -1.47 | 892 | *Mag* |
| HP_ONSET | 0.00016 | 0.045 | -0.32 | -1.42 | 846 | *Mag* |
| MEISSNER_BRAIN_HCP_WITH_H3K4ME3_AND_H3K27ME3 | 0.00016 | 0.045 | -0.41 | -1.77 | 829 | *Cldn11* |
| GOBP_CENTRAL_NERVOUS_SYSTEM_DEVELOPMENT | 0.00016 | 0.045 | -0.41 | -1.77 | 758 | Mag |
| FAN_EMBRYONIC_CTX_OLIG | 0.00016 | 0.045 | -0.51 | -2.19 | 614 | Cldn11, Mag |
| GOBP_NERVOUS_SYSTEM_PROCESS | 0.00016 | 0.045 | -0.34 | -1.46 | 655 | Mag, Col11a1 |
| HP_CONSTITUTIONAL_SYMPTOM | 0.00016 | 0.045 | -0.34 | -1.46 | 643 | Col11a1 |
| Oligodendrocyte_All_Zeisel_Science_2015 | 0.00017 | 0.045 | -0.59 | -2.40 | 320 | Cldn11, Mag |
| GOBP_GLIOGENESIS | 0.00017 | 0.045 | -0.51 | -1.99 | 218 | Mag |
| GOBP_GLIAL_CELL_DIFFERENTIATION | 0.00018 | 0.045 | -0.55 | -2.09 | 168 | Mag |
| DESCARTES_MAIN_FETAL_SCHWANN_CELLS | 0.00018 | 0.045 | -0.56 | -2.10 | 159 | Mag |
| ZHONG_PFC_C4_PTGDS_POS_OPC | 0.00018 | 0.045 | -0.52 | -1.88 | 120 | Cldn11, Mag |
| GOBP_ENSHEATHMENT_OF_NEURONS | 0.00018 | 0.045 | -0.60 | -2.12 | 105 | Cldn11, Mag |
| GOBP_GLIAL_CELL_DEVELOPMENT | 0.00019 | 0.045 | -0.60 | -2.05 | 89 | Mag |
| GOBP_OLIGODENDROCYTE_DIFFERENTIATION | 0.00019 | 0.045 | -0.63 | -2.10 | 78 | Mag |
| LEIN_OLIGODENDROCYTE_MARKERS | 0.00019 | 0.045 | -0.82 | -2.69 | 70 | Mag |
| DESCARTES_MAIN_FETAL_OLIGODENDROCYTES | 0.00019 | 0.045 | -0.69 | -2.19 | 59 | Cldn11, Mag, Col11a1 |
| GOCC_MYELIN_SHEATH | 0.00019 | 0.045 | -0.75 | -2.17 | 35 | Mag |
| Oligodendrocyte_Myelinating_Zhang_JNeuro_2014 | 0.00019 | 0.045 | -0.85 | -2.47 | 35 | Cldn11 |
| Oligodendrocyte_All_Cahoy_JNeuro_2008 | 0.00019 | 0.045 | -0.87 | -2.48 | 33 | Cldn11, Mag |
| DESCARTES_FETAL_CEREBRUM_OLIGODENDROCYTES | 0.00019 | 0.045 | -0.76 | -2.18 | 33 | Cldn11, Mag |
| Oligodendrocyte_Mature_Darmanis_PNAS_2015 | 0.00020 | 0.045 | -0.91 | -2.25 | 17 | Cldn11, Mag |

## Discussion

Our research provides novel insight into the effect of ELS on the PFC. ELS is known to cause a profound and lasting impact on the brain, leading to cognitive impairments, heightened sensitivity to future stressors, and increased risk for mental health disorders. Therefore, studying ELS in the PFC is important because the PFC plays a key role in cognitive functions like decision-making, impulse control, emotional regulation, and social behavior (Arnsten, 2009). During childhood and adolescence, the PFC undergoes significant development and experiences increased vulnerability to stress (Chocyk et al., 2013). When stress occurs during this critical period, it can alter the development of neural circuits within the PFC, heightening the risk of mental health disorders such as depression, anxiety, and PTSD (Colich et al.,2017). Additionally, ELS affects neuroplasticity—the brain’s ability to adapt and reorganize—resulting in impairments in learning, memory, and other executive functions (Malave et al., 2022).

To gain deeper insights into how ELS influences the PFC, we conducted a meta-analysis of publicly available transcriptional profiling datasets. We identified five relevant datasets (GSE89692, GSE116416, GSE14720, GSE153043, GSE124387) that examined the long-term effects of multi-day ELS paradigms, such as maternal separation for >3 hours per day and/or limited nesting material on male and female laboratory pups (rats, mice) during the postnatal period (between postnatal days P1-P21). The outcome variable was gene expression in the PFC later in life (in adulthood) as measured by microarray or RNA-Seq. We achieved stable meta-analysis estimates for 11,885 genes, identifying five key results that withstood false discovery rate correction (FDR<0.05): the down-regulation of *Tgfa, Cldn11, Iqgap3, Col11a1*, and *Mag*.

Notably, the proteins encoded by *Mag* and *Cldn11* are both important for myelination. *Mag* encodes a membrane protein that is part of the immunoglobulin superfamily and is thought to be involved in the maintenance of normal axon myelination and some neuron-myelin communication (Quarles, 2007). *Cldn11* produces a protein that is part of the Claudin family, which is a crucial component of tight junctions in cell membranes. Tight junctions act as barriers, preventing the free passage of substances between cells and maintaining cell structure and communication. Claudin 11 is a myelin component and an important regulator of oligodendrocyte proliferation and migration (Riedhammer et al., 2021). In addition to *Mag* and *Cldn11, Tgfa* is another down-regulated gene that survived false discovery rate correction and plays a protective role in maintaining oligodendrocyte viability and white matter integrity, as demonstrated in models of cerebral ischemia where its deficiency led to exacerbated cell death in oligodendrocytes and impaired myelin maintenance (Dai et al., 2020).

When conducting a gene set enrichment analysis to explore the functional patterns within our results, we identified 42 gene sets that were enriched with the effects of ELS, the majority of which were down-regulated and related to brain development, especially oligodendrocyte development and myelination, lead by down-regulation of *Cldn11, Mag*, and *Col11a1*. We also observed an enrichment of upregulation of genes that had previously been shown to be upregulated in the cerebral cortex of patients with Major Depressive Disorder. However, the strongest upregulated gene in this set, *Fst* —the product of which is an inhibitor of follicle-stimulating hormone release - did not surpass our threshold for differential expression (FDR>0.10) (Shi et al., 2016). The StringDB database as well revealed that the protein products of the down-regulated genes *Cldn11* and *Mag* are part of a predicted PPI network with the protein products for other genes that showed trends towards significant differential expression (FDR<0.10) in association with ELS in our meta-analysis, including down-regulated *Mal, T Cell Differentiation Protein* (*Mal*), also known as *Myelin And Lymphocyte Protein, Kallikrein related-peptidase 6* (*Klk6)*, and *Fatty acid 2-hydroxylase (Fa2h)*, and upregulated *UDP-glucose ceramide glucosyltransferase* (*Ugcg)* (Szklarczyk et al., 2023). According to StringDB, this network is enriched in genes associated with oligodendrocyte differentiation and myelination, including *Mag, Mal, Klk6, Fa2h*, and *Ugcg*. Collectively, these findings suggest that ELS during critical developmental periods may lead to long-term disruptions in myelin-related processes, potentially affecting the efficiency of neural transmission in the PFC.

Altogether, our meta-analysis supports previous evidence in rodents, demonstrating that ELS disrupts the normal development and differentiation of oligodendrocytes and the process of myelination, essential components of cortical white matter. For instance, Yang et al. (2017) demonstrated that separating rat pups from their mothers for 3 hours daily during the first three weeks of life reduces myelination in the PFC (Yang et al., 2017). This reduction in myelination was linked to a decrease in mature oligodendrocytes and an increase in oligodendrocyte precursor cells (OPCs), suggesting that ELS impairs normal maturation, preventing OPCs from properly differentiating into mature myelinating cells. These myelination deficits were observed both after weaning (at P21) and in adult rats (at P60). Likewise, Teissier et al. (2020) found that when mice were exposed to 3 hours of split litter maternal separation (slMS) during the first two weeks of life, it caused early maturation of OPCs in the PFC by day 21 (P21), followed by their depletion in adulthood (Teissier et al., 2020). Makinodan et al. (2012), using PLP-eGFP mice, discovered that isolating juvenile mice after weaning (P21-35) led to abnormal maturation of oligodendrocytes in the PFC and impaired performance on cognitive tasks (Makinodan et al., 2012).

Further, Abraham et al. found that early life adversities seem to impact the development of white matter, particularly affecting oligodendrocytes. These changes in myelination were observed in brain regions that are maturing during the period when these early adversities occur (Abraham et al., 2023). Of note, exposure of adult rodents to chronic social stress has also been shown to impair myelination in the PFC (Bonnefil et al., 2019; Liu et al., 2012).

Preventing disruptions in oligodendrocyte maturation can mitigate some long-term behavioral effects of ELS. For example, Yang et al. (2017) obtained evidence suggesting that maternal separation leads to a long-term increase in WNT signaling, which blocks OPC differentiation and new myelination in the adult PFC (Yang et al., 2017). Importantly, administering a WNT antagonist (XAV939) during the first three weeks of life partially rescued the myelination and behavioral deficits observed after ELS. This study is significant as it is the first to demonstrate that agents that reverse myelination deficits can rescue behavioral deficits in a rat model of early adversity. Additionally, Teissier et al. (2020) explored the impact of neuronal activity in the developing PFC on myelination and found that inhibiting neuronal activity during the first two weeks of life led to early myelination and similar behavioral problems as seen in mice exposed to maternal separation (Teissier et al., 2020). Conversely, activating PFC neurons from days 2 to 14 resulted in reduced myelination at P15 and increased myelination in adulthood, which reversed the behavioral deficits observed in these models.

ELS has been linked to altered oligodendrocyte development and myelination in children, showing that these changes extend beyond animal models to human development. Hanson et al. (2013) examined white matter properties and neurocognitive performance in children who experienced early neglect compared to those raised in typical environments (n = 63, average age = 11.75 years). Their findings revealed that children who experienced early neglect displayed alterations in prefrontal white matter microstructure, characterized by a more diffuse organization, which was associated with neurocognitive deficits. This indicates that ELS-induced abnormalities in white matter can contribute to cognitive deficits, particularly in the PFC, as a result of neglect (Hanson et al., 2013).

Our meta-analysis both supports and extends these previous findings, offering a glimpse at the molecular changes in the PFC in response to ELS that may underlie myelination deficits and persist into adulthood. By using a meta-analysis to study the effect of ELS on the PFC, we were able to combine data from multiple studies to effectively reach a final sample size of 89, increasing statistical power to a level capable of detecting moderate effect sizes and providing more reliable estimates of the effects.

Furthermore, by including studies from multiple ELS models (maternal separation, limited bedding and nesting) and laboratory settings in our meta-analysis, we identified consistent patterns and enhanced the generalizability of our findings. However, the we did not have the power to identify patterns that were ELS model specific, as only five final studies survived our inclusion/exclusion criteria for the meta-analysis. This was partially due to our high standard for the quality of the included studies – for example, we ended up rejecting one dataset that lacked an associated publication (GSE23728), preventing us from fully reviewing its methodology. To maintain consistency in data processing, we also only extracted preprocessed transcriptional profiling datasets from the Gemma database. As future datasets related to the effects of ELS on the PFC are released, updating our meta-analysis will likely result in additional significant findings.

## Conclusion

Approximately 65% of people in the United States experienced at least one type of ELS, while 12.5% experienced as many as four types (Fogelman & Canli, 2019). Characterizing the impact of ELS on the PFC is crucial for understanding the long-term cognitive and behavioral effects of this early stress. Our results suggest that ELS during critical periods of development may produce long-term effects on molecular contributors to oligodendrocyte development and myelination, potentially altering the efficiency of transmission in the PFC. Our results also indicate that ELS can drive gene expression changes similar to those observed in the frontal cortex of patients with Major Depressive Disorder. This knowledge is vital for developing targeted interventions aimed at reducing the negative consequences of ELS and improving mental health outcomes.

## Acknowledgements

This work was supported by the Hope for Depression Research Foundation (HDRF: HA, MJM, EJN), Grinnell College Center for Careers, Life, and Service (TQD & AB), the International Brain Research Organization (IBRO) and Faculty for Undergraduate Neuroscience (FUN) (TQD & DMN), NIDA (U01 DA043098: HA), the Pritzker Neuropsychiatric Disorders Research Foundation (HA & SJW). Funders and sponsors had no active role in the review.

