## Supplemental Table Legends & Figures for "A meta-analysis of the effects of early life stress on the prefrontal cortex transcriptome suggests long-term effects on myelin"

***Table S1. The full meta-analysis results (11,889 genes, 11,885 stable meta-analysis estimates).*** *This .xlsx file includes two worksheets: 1) The worksheet "MetaAnalysisOutputByPval" provides the full meta-analysis results, with each row representing the results for one gene, and each column providing either gene annotation or meta-analysis statistical output. The results are ordered by p-value, so that the top rows in the worksheet are the genes with the smallest p-values. 2) The worksheet "ColumnDefinitions" provides the definitions for the variables present in each column in "MetaAnalysisOutputByPval".*

***Table S2: The full fast Gene Set Enrichment Analysis (fGSEA) results (9322 gene sets).*** *This .xlsx file includes two worksheets: 1) The worksheet "fGSEA_Results" provides the full fGSEA results, with each row representing the results for one gene set, and each column providing the fGSEA statistical output. The results are ordered by p-value, so that the top rows in the worksheet are the gene sets with the smallest p-values. 2) The worksheet "ColumnDefinitions" provides the definitions for the variables present in each column in "fGSEA_Results".*

**Supplementary Figures**

**A.**

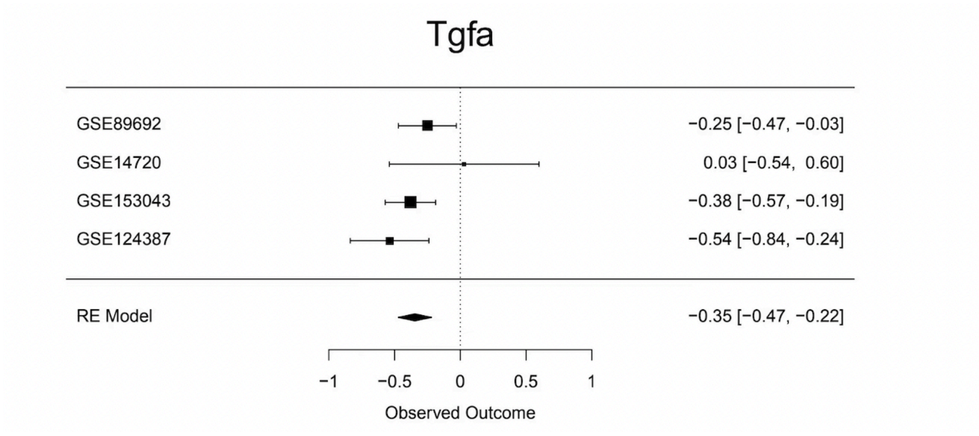


**B.**

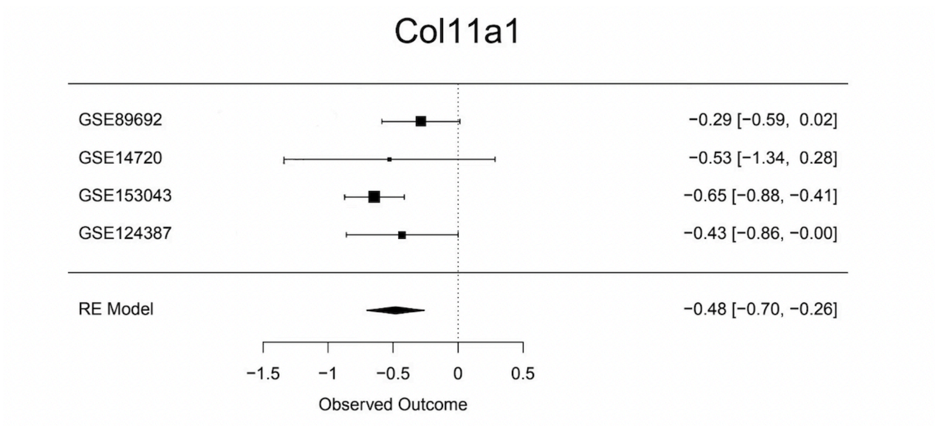


**C.**

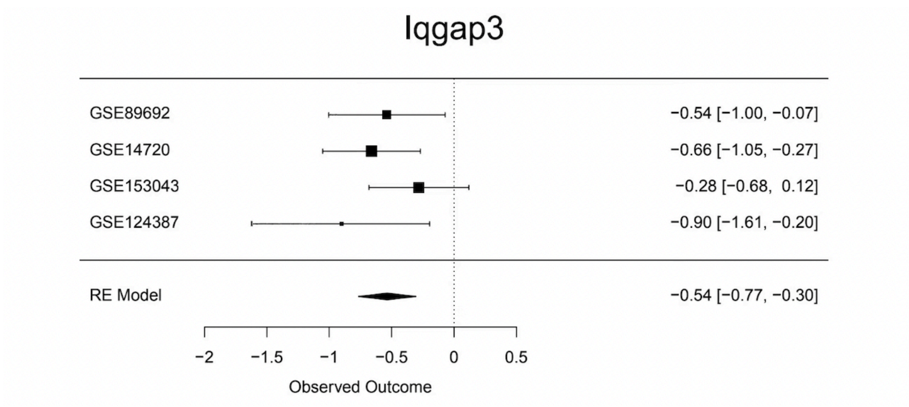


***Figure S1. Forest plots illustrating the differential expression following ELS. A-C.*** *Forest plots for the three differentially expressed genes from the meta-analysis (FDR<0.05) that weren't illustrated in the main text. Rows illustrate ELS Log2FC (squares) with 95% confidence intervals (whiskers) for each of the datasets and the meta-analysis random effects model ("RE Model"). Forest plots allow for visual inspection of the consistency and magnitude of effects across the five studies.* ***A.*** *A forest plot showing the*

*down-regulation of transforming growth factor alpha (Tgfa) in ELS models.* ***B.*** *A forest plot showing the down-regulation of collagen, type XI, alpha 1 (Col11a1) in ELS models.* ***C.*** *A forest plot showing the down-regulation of IQ motif containing GTPase activating protein 3 (Iqgap3) in early life stress models.*
